# Split-Indigoidine synthetase as optical reporter for benchmarking protein-protein interactions

**DOI:** 10.64898/2026.06.03.729908

**Authors:** Patrick Gonschorek, Christian Schelhas, Melina Flakowski, Leonie Schenk, Adrian Podolski, Helge B. Bode

## Abstract

Indigoidine is a blue pigment biosynthesized by a single-module Non-Ribosomal Peptide Synthetase (NRPS) using L-glutamine as substrate. Despite its potential as a colorimetric reporter, no such system has been established from it to date. We used a recently characterized interdomain fusion site located between its adenylation (A) and thiolation (T) domains to develop the Indi2GO system, which provides a naked-eye detectable and quantitative optical readout of transient and covalent protein-protein-interaction (PPI) in living cells. Indi2GO enables high-throughput benchmarking and optimization of PPI tools in a standard 96-well plate reader format, without requiring exogenous substrates, specialized equipment or complex analytical workflows. We demonstrate its broad applicability with three widely used protein-protein interaction tools: SYNZIPS, inteins, and the SpyTag:SpyCatcher system. We used Indi2GO to validate novel SYNZIP pairs, which we used in NRPS engineering, highlighting its applicability for the development of novel PPI-mediating tools in the context of NRPS engineering and synthetic biology.

## INTRODUCTION

Indigoidine is a blue-pigmented natural product synthesized by bacteria such as *Pseudomonas, Streptomyces* and *Photorhabdus* via indigoidine synthetases^1^. It is thought to protect its producers against oxidative stress by capturing reactive oxygen species^2^.

Indigoidine synthetases (IndC or BpsA) are Non-Ribosomal Peptide Synthetases (NRPS), which generally function via a thiotemplate mechanism, whereby substrate amino acids are activated by adenylation (A) domains, covalently tethered to thiolation (T) domains, connected by condensation (C) domains, and ultimately released by a thioesterase (TE) domain. NRPS enzymes are initially produced in an inactive *apo*-form and require post-translational activation by a phosphopantetheinyl transferase (PPTase), which transfers the phosphopantetheinyl group from coenzyme A (CoA) to a conserved serine in the T domain to generate the active *holo*-enzyme^3^.

In contrast to the multimodular complexity of most NRPS assembly lines, BpsA/IndC are single-module enzymes comprising only one A, T, and TE domain each, and additionally, embedded within the A_sub_ domain, an oxidase (Ox) domain. First, glutamine is activated as aminoacyl adenylate by the A domain, requiring ATP, following its covalent binding onto the T domain as thioester. The T domain bound L-glutamine is then oxidized by the Ox domain, using flavin mononucleotide (FMN) as electron acceptor before the oxidized glutamine is finally cyclized by the TE domain and released. In the presence of oxygen, two monomers dimerize to form the blue pigment indigoidine^4^.

Due to these rather simple requirements, IndC/BpsA have previously been explored as reporters for screeninging inhibitors of PPTase activity^5^, as a reporter for genome editing^6^ and to boost natural product production in *Streptomyces*^7^, as visual whole-cell biosensor for heavy metals^8^, and as a reporter for the production of an non-ribosomal peptide in mammalian cells^9^. Furthermore, the enzyme has been engineered for enhanced catalytic activity using its own pigment production as readout^10^.

Split-protein variants of many optical reporters have been developed to detect protein-protein interactions (PPIs), whereby the reporter protein is separated into two inactive halves that are reconstituted by PPI-mediated complementation. Prominent examples include bimolecular fluorescence complementation (BiFC)^11,12^, split-luciferases^13,14^, and split-enzymes converting chromogenic substrates^15^, each offering distinct advantages in terms of sensitivity, dynamic range, and spatial and temporal resolution. Notably, while split-fluorescent proteins function without exogenous substrates, split enzymes require externally supplied cofactors such as furimazine for split-NanoLuc or 5-bromo-4-chloro-indolyl-galactopyranoside (X-gal) for split-β-galactosidase. To our knowledge, no split chromogenic reporter system operating without exogenous substrate addition has been reported to date.

Here we describe **Indi2GO** (**Indi**goidine-based **G**enetic **O**ptical readout), a novel colorimetric split-reporter system for the quantification of PPIs in the context of synthetic biology and NRPS engineering. By splitting the indigoidine NRPS at its XUT^I^ site, we generate a quantitative PPI reporter that operates in a standard *E. coli* DH10B::*mtaA* strain and is fully compatible with 96-well high-throughput screening formats. We demonstrate the utility of Indi2GO by systematically benchmarking the three widely used PPI tools SYNZIPS^16^, inteins^17^, and the SpyTag:SpyCatcher^18^ system resolving differences in interaction strength. Uniquely among split chromogenic reporters, Indi2GO requires no exogenous substrate, exploiting instead the intracellular L-glutamine pool to convert the strength of PPIs into visible blue signals.

## RESULTS & DISCUSSION

### Splitting the indigoidine synthetase

To assess the potential of the indigoidine as colorimetric output of PPIs, we took the indigoidine sythetase encoding *indC* from *Photorhabdus laumondii* TT01^19^ and expressed it heterologusly in the *E. coli* DH10B::*mtaA* strain, which constitutively expresses a broad-spectrum PPTase. IndC expression yielded robust pigment production, in contrast to the uninduced culture (**Fig. 1a**). Pigment production was quantified spectrophotometrically by measuring OD_600_, corrected for biomass-contribution using OD_800_ of a reference culture, as described previously^20^ and described in the supplementary material (**Supplementary Fig. S1**). Pigment concentrations peaked at 30 h post-induction and remained stable through 48 h, with only a minimal decline (**Fig. 1b**), demonstrating that a single absorbance measurement after 48 h is sufficient to quantify pigment production without the need for kinetic measurements. This stability is consistent with previous reports that indigoidine can be maintained under standard aerobic conditions, although gradual bleaching may occur over prolonged timeframes due to its reactive oxygen species scavenging activity; supplementation with ascorbic acid has been reported to prevent this degradation, although this was not employed in the present study. Notably, treatment with reducing agents converts the blue-colored keto form of indigoidine into its colorless leuco isoform, which exhibits excitation at 415 nm, rendering it amenable to fluorescence microscopy applications and further expanding the analytical utility of the Indi2GO platform^9,21^.

**Figure 1.**
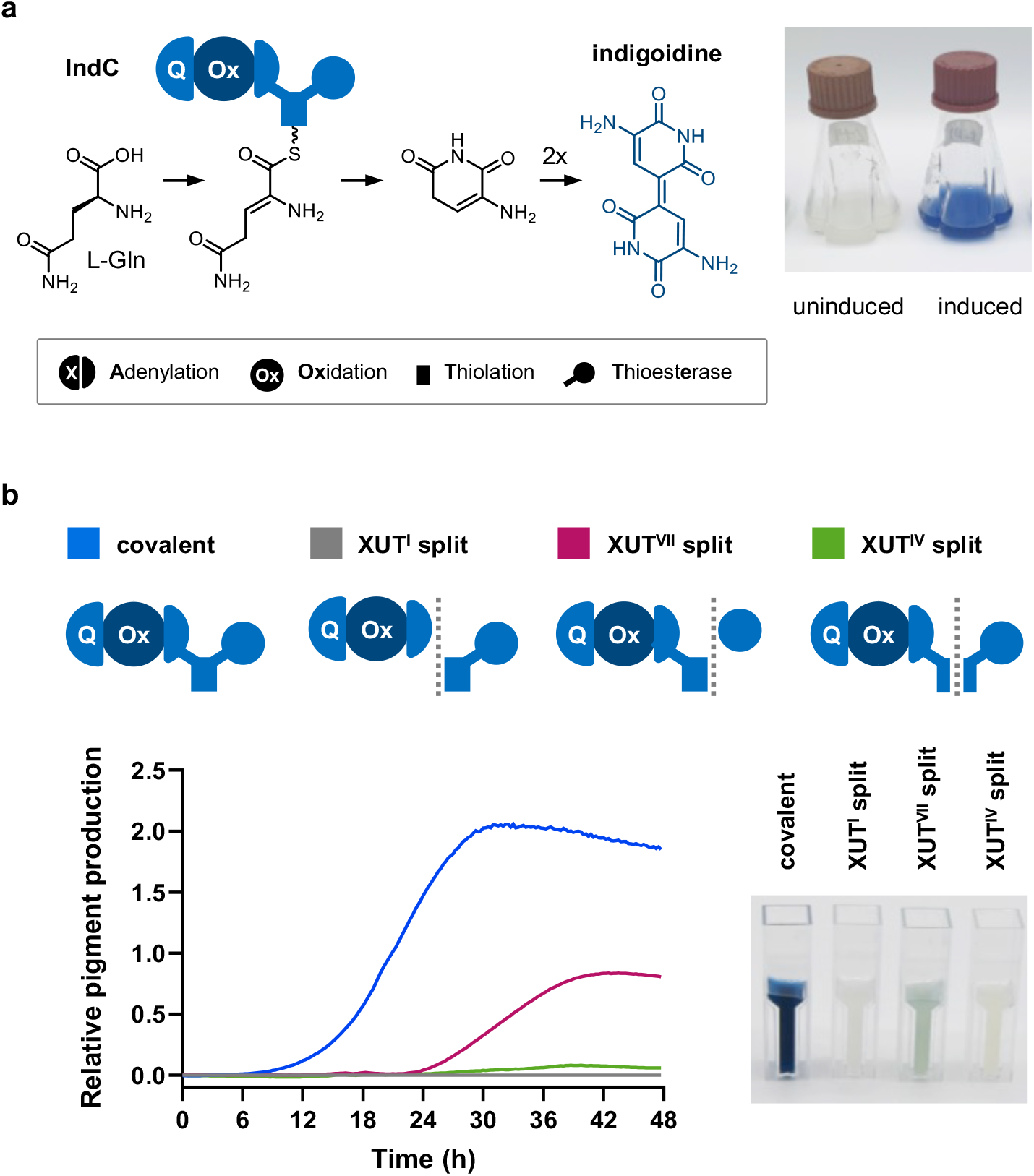
Identification of suitable split sites within the indigoidine synthetase IndC. (**a**) Schematic representation of the IndC domain architecture (A, Ox, T, TE domains) and indigoidine biosynthesis. (**b**) Three XUT split sites (XUT^I^, XUT^IV^, XUT^VII^) and time-course of relative indigoidine production of full-length IndC and split variants over 48 h post-induction. Splitting at XUT^I^ and XUT^IV^ completely abolished indigoidine production, while splitting at XUTV^II^ retained approximately 44% of full-length activity.

Next, we sought to identify a split site that would completely abolish pigment production without spontaneous self-assembly of the resulting fragments, which would cause background signal. This was particularly relevant in the context of IndC given that a previous study of the indigoidine synthetase BpsA from *Streptomyces lavendulae* reported catalytic activity when the enzyme was split between T and TE domain^22^. We relied on the recently defined fusion sites between A and T domain termed “eXchange Units between T domains” (XUT)^23^, providing three rational split points: one upstream (XUT^I^), one in the middle (XUT^IV^) and one downstream (XUT^VII^) of the T domain. We then split the *indC* gene at each site and expressed the resulting fragments from two separate plasmids^22^.

Interestingly, splitting at XUT^VII^, which corresponds to the previously described BpsAΔTE – TE split^22^, retained 44% of the native IndC pigment production. In contrast, splitting at either XUT^I^ or XUT^IV^ completely abolished pigment production (**Fig. 1b**). Therefore, both XUT^I^ and XUT^IV^ would be suitable split sites for reporter development. We selected XUT^I^ over XUT^IV^ based on previous findings that demonstrated a greater tolerance for insertions at the interdomain XUT^I^ site compared to the intradomain XUT^IV^ site^24^.

### Reconstituting the split indigoidine synthetase with PPI-mediating tools

After having identified a split site that abolishes pigment production, we though to restore it using PPI-mediated complementation. To evaluate the potential of our split reporter across a broad spectrum of interaction types, we selected PPI tools spanning both non-covalent and covalent interactions of varying strength. For covalent interactions, we included two mechanistically distinct systems: (i) The split intein NrdJ-1, which autocatalytically excises itself leaving only a six amino acids scar sequence within the A-T linker^24^. (ii) The SpyTag:SpyCatcher 003 system, which forms a covalent iso-peptide bond between a lysine on SpyCatcher and an aspartate on SpyTag, resulting in a covalent complex which remains at the protein-protein interface^18^. For non-covalent interactions, we selected two synthetic zipper (SYNZIP, SZ) pairs with different orientation: The parallel pair SZ1:SZ2 and the antiparallel pair SZ17:SZ18 (**Supplementary Fig. S2**)^16,25^. PPI-mediated complementation of the split IndC fragments successfully restored blue pigment production across all tested interaction tools to varying levels (**Fig. 2**). The covalent SpyCatcher:SpyTag system and the split intein NrdJ-1 restored pigment production to 104 and 78% of native IndC levels, respectively. Striking differences were observed among the SYNZIP pairs tested: While the parallel pair restored indigoidine production to only 9%, the antiparallel pair restored to 101% of the native pigment production. These results are consistent with our previous observations that antiparallel SZs, which position the interacting C- and N-termini in close proximity, are inherently better suited for split-protein complementation than parallel SZs, in which the termini are oriented away from each other, generating a gap between the two protein fragments^26^. Collectively, we validate split-IndC complementation successfully with three mechanistically distinct interaction tools.

**Figure 2.**
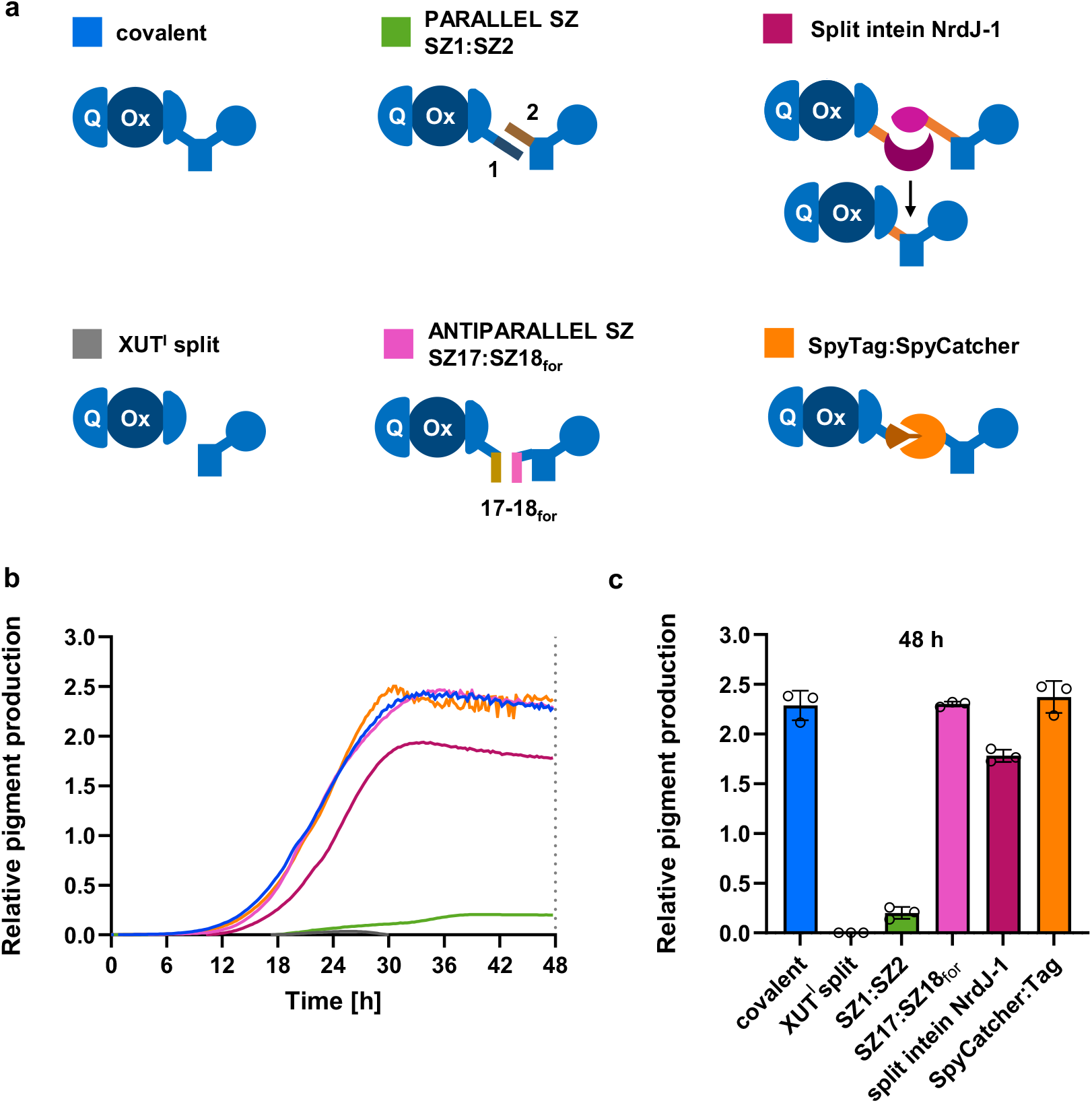
Insertion of protein-protein interaction tools into the split indigoidine synthetase. (**a)** Schematic of the split-IndC construct architecture with the respective PPI tools inserted at the XUT^I^ site. (**b**) Time-course of relative indigoidine production of split-IndC constructs carrying different PPI tools over 48 h post-induction. (**c**) Relative indigoidine production at 48 h Data represent mean ± SD of biological triplicates (n = 3). Wild-type IndC synthetase served as a positive control; the unfused XUT^I^ split variant served as a negative control. Parallel SYNZIP pairs (SZ1:SZ2) restored only minimal indigoidine production, whereas antiparallel configurations based on SZ17:SZ18 (SZ17:SZ18for and SZ17:SZ18rev) restored indigoidine production to levels comparable to wild-type IndC synthetase. The split intein and the SpyTag:SpyCatcher003 system similarly enabled efficient restoration of catalytic activity.

To further optimize the antiparallel SZ17:SZ18-based pairs, we systematically evaluated the effect of inserting one to three GSG linker sequences between the SYNZIPs and split-IndC fragments. In both cases, the insertion of a single GSG linker unit increased pigment production to more than 120% of native IndC levels (**Supplementary Fig. S3**).

### Assessing the dynamic range of Indi2GO using truncated SYNZIPs

A broadly applicable PPI reporter should not only provide binary interaction readouts but also resolve differences in binding affinity. Stepwise truncation of SYNZIPs enables systematic modulation of binding affinity while preserving both the interaction partners and the spatial distance of the fused protein fragments. To this end, we selected the antiparallel SYNZIP pair SZ17:SZ18 (42 and 41 amino acids, respectively), which exhibits a reported dissociation constant of less than 10 nM ^25^, as our model system. To systematically modulate interaction affinity of the SYNZIPS, we designed a sequential truncation series in which both SZ17 and SZ18 were shortened by 10 amino acids in successive steps from their distal, freestanding termini (C-terminus for SZ17, N-terminus for SZ18), thereby preserving the spatial relationship between the two IndC fragments (**Fig. 3a, d**).

**Figure 3.**
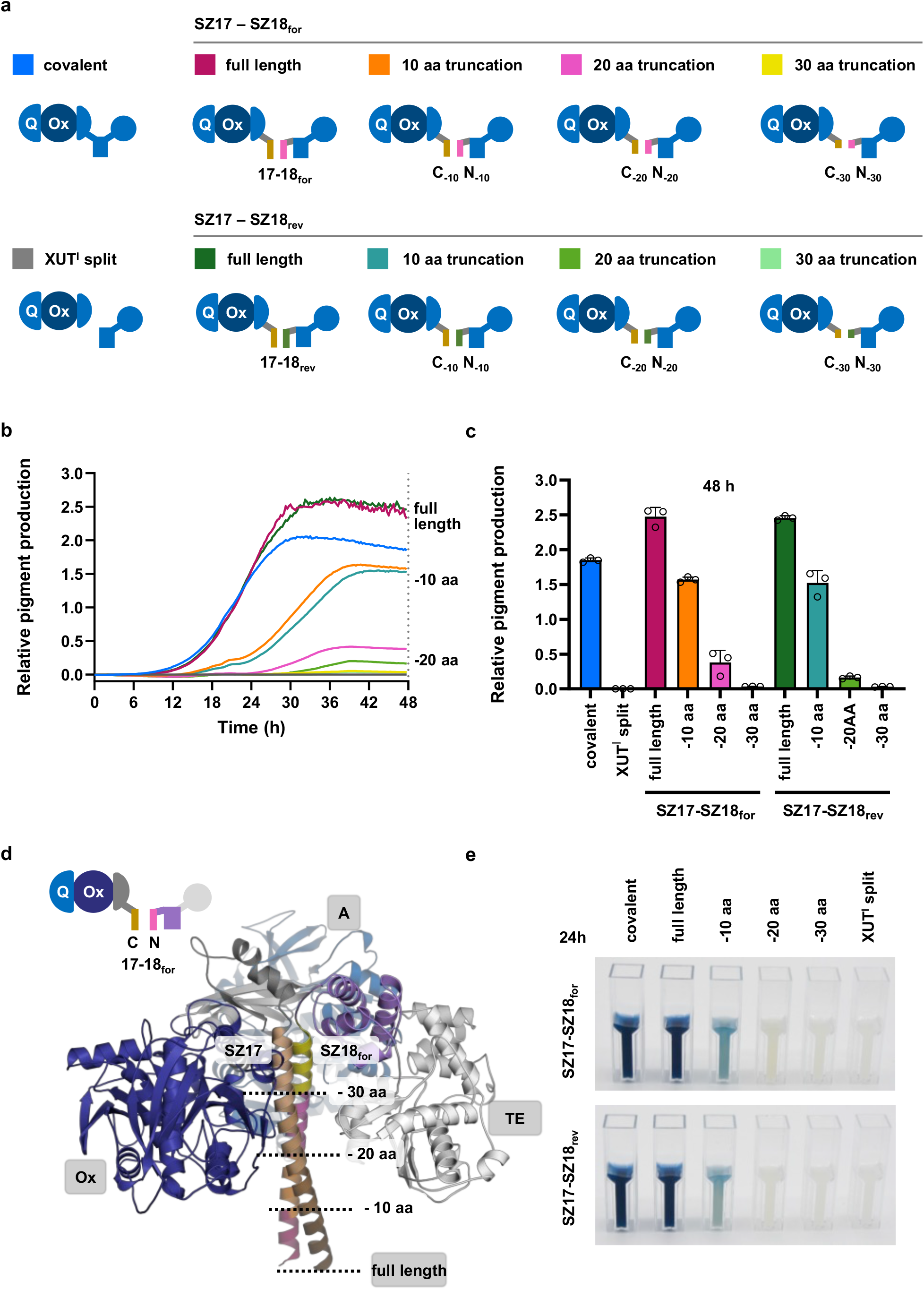
Truncating SYNZIPs to assess dynamic range of the system. (**a**) Schematic of split-IndC with progressively truncated SZ17:SZ18 variants, shortened stepwise from the distal ends of both SYNZIPs. (**b**) Time-course of relative indigoidine production of split-IndC constructs carrying full-length and truncated SZ17:SZ18for and SZ17:SZ18rev pairs over 48 h post-induction. (**c**) Relative indigoidine production at 48 h. Mean ± SD of biological triplicates (n = 3). (**d**) AlphaFold 3 structural model of the SZ17:SZ18for-fused split-IndC construct, with dashed lines indicating the successive truncation points applied to both SYNZIP partners. (**e**) Representative culture image showing chromogenic output across full-length and truncated SYNZIPs.

While conducting this truncation series, we made the observation that the sequence of SZ18 had been used in the reverse orientation (C-to N-terminus) in previous works^27-29^, effectively generating an inverted variant, hereafter referred to as SZ18_rev_, while the original sequence as described in the initial publication^16^, is referred to as SZ18_for_. Interestingly, this inversion seemed to have had no obvious impact on its performance in previous works. To investigate the impact of this sequence inversion of SZ18 on affinity and orientation, we compared both variants in the Indi2GO system and conducted the truncation series with both variant of SZ18, SZ18_for_ and SZ18_rev_.

To visualize the truncation series, AlphaFold 3 models of the corresponding constructs were generated, illustrating the progressive reduction in coiled-coil length with each truncation step (**Fig. 3d**). Within the Indi2GO system, both full-length SZ17:SZ18 pairs sustained indigoidine production at approximately 134% of native IndC levels (**Fig. 3c**). Truncation of SZ17:SZ18_for_ by 10 amino acids caused a significant reduction in pigment yield to 85% of native IndC levels after 48 h. Truncation by 20 amino acids decreased production to 21%, while removal of 30 amino acids nearly abolished pigment formation, reducing yield to 2% of native IndC levels. Strikingly, SZ17:SZ18_rev_ behaved similar in the truncation series: While full-length variant sustained indigoidine production at 133% truncation by 10 amino acids reduced pigment yield to 82%, while truncation by 20 amino acids resulted in a more pronounced drop to 9% of native IndC levels. As observed for SZ17:SZ18_for_, removal of 30 amino acids nearly abolished pigment formation, reducing yield to 2% of native IndC levels.

Time-course analysis revealed that diminished interaction not only result in a lower final figment yield, but in lower production rates resulting in a delayed timepoint of reaching this maximum (**Fig. 3b**). Importantly, endpoint measurements seem to correlate with PPI affinity (**Fig. 3c**), and the gradual chromogenic output was visible by the naked eye without the need for analytical instrumentation (**Fig. 3e**). Collectively, these data establish that Indi2GO provides a quantitative, affinity-dependent colorimetric readout over a meaningful dynamic range, rendering it suitable for the comparative analysis of PPI strengths.

### Probing split-protein fragment distance through directional SYNZIP inversion

Based on the observation that parallel and antiparallel SYNZIP behave fundamentally different, we hypothesized that distance between the split-IndC fragments at the XUT^I^ site also influences pigment productions. Building on the observation that certain SYNZIP sequences retain functionality upon inversion, we comprehensively investigated whether such inversion could convert parallel SYNZIP pairs into antiparallel orientation, thereby bringing the split-IndC fragments into closer proximity.

SYNZIPs were originally designed to enforce interactions in a defined orientation, however the molecular consequences of inverting one SYNZIPs sequence are difficult to predict a priori. Therefore, we generated AlphaFold 3 models of SYNZIP pairs and measured the predicted distances between the residues at the XUT^I^ split site (**Supplementary Fig. S4**). As a benchmark, AlphaFold 3 accurately reproduced the known crystal structures of SZ5_for_:SZ6_for_ (**Supplementary Fig. S4a**, SZ5_for_:SZ6_for_, PDB ID: 3HE4 displayed in grey) and SZ1_for_:SZ2_for_ (**Supplementary Fig. S4b**, SZ1_for_:SZ2_for_, PDB ID: 3HE5 displayed in grey). AlphaFold3 structural predictions supported the proposed hypothesis that inverting a single SYNZIP sequence lead to an inversion of orientation form parallel to antiparallel while simultaneous inversion of both partners restored the original orientation.

To experimentally test the effects of SYNZIP direction and its impact on protein fragment distance, we evaluated all four orientation combinations of the two SYNZIP pairs SZ1:SZ2 and SZ5:SZ6 in the Indi2GO system. Notably, both pairs have experimentally validated affinities in the same range (< 10 nM for SZ1_for_:SZ2_for_ and < 15 nM for SZ5_for_:SZ6_for_)^25^.

For both SYNZIP pairs tested, parallel configurations consistently yielded large predicted interdomain distances and poor indigoidine production: SZ1_for_:SZ2_for_ and SZ1_rev_:SZ2_rev_ produced only 9 and 0% of native indigoidine levels at distances of 72.9 Å and 69.5 Å, respectively, while both parallel SZ5:SZ6 configurations (59.0 Å and 65.5 Å) completely abolished indigoidine production. In contrast, all antiparallel configurations displayed substantially shorter predicted interdomain distances and correspondingly higher indigoidine yields, ranging from 32 to 84% of native indigoidine levels at distances between 13.8 Å and 30.4 Å (**Fig. 4c**). Across both SYNZIP pairs, indigoidine production correlated inversely with predicted interdomain distance, consistently demonstrating that sequence inversion of one SYNZIP partner converts parallel pairs into antiparallel configurations with substantially improved reconstitution efficiency.

**Figure 4.**
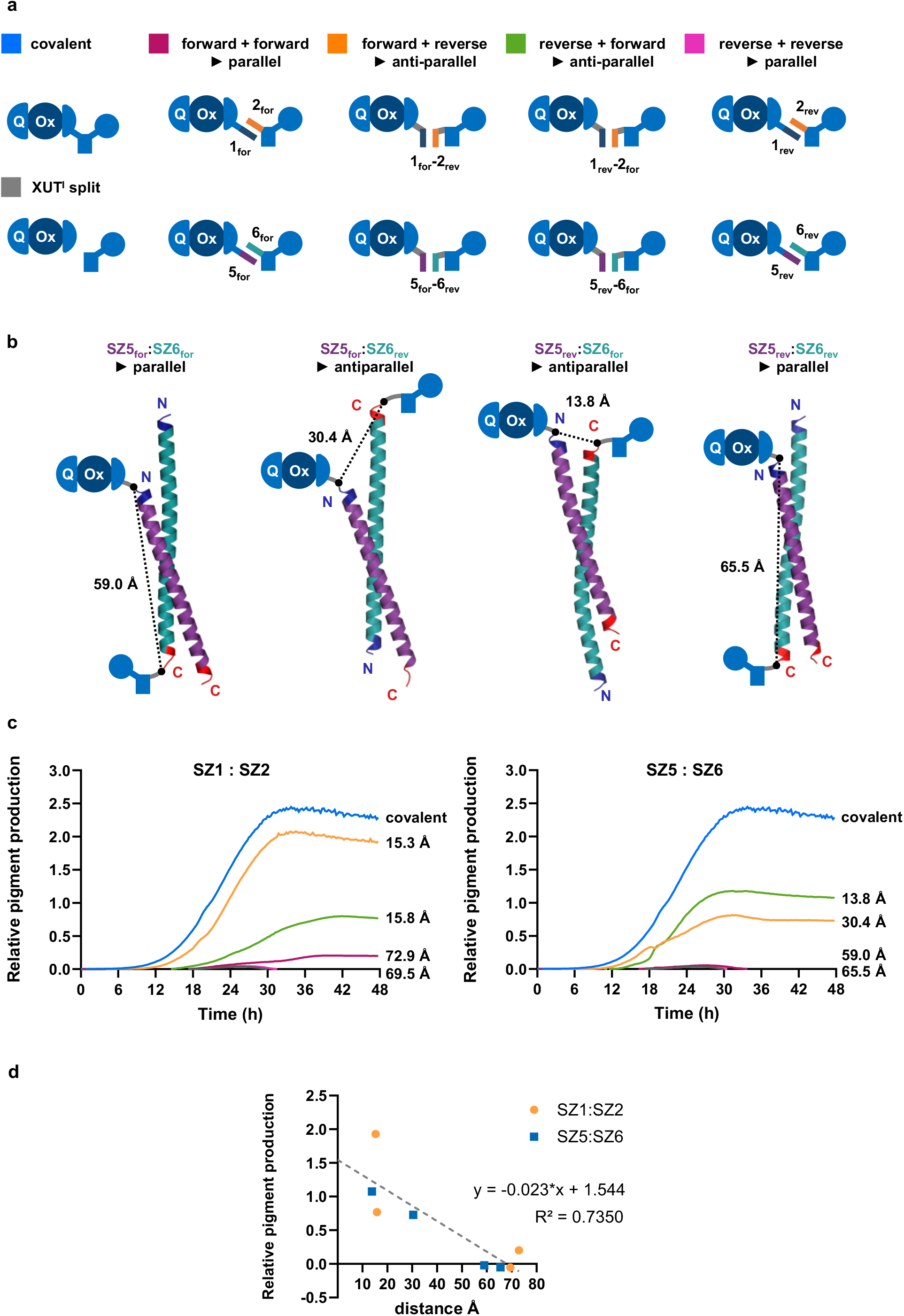
Development of novel antiparallel SYNZIPs using the Indi2GO system. (**a**) Schematic representation of the split-IndC construct with all four orientation combinations of SYNZIPs SZ1:SZ2 and SZ5:SZ6 inserted at the XUT^I^ site, illustrating the parallel (SZ1for:SZ2for, SZ1rev:SZ2rev) and antiparallel (SZ1for:SZ2rev, SZ1rev:SZ2for) configurations. (**b**) AlphaFold 3 structural models of all four orientation combinations for SZ1:SZ2 and SZ5:SZ6, with the predicted interdomain distance. (**c**) Time-course of relative indigoidine production for split-IndC constructs with all four orientation combinations of SZ1:SZ2 and SZ5:SZ6 over 48 h post-induction; predicted interdomain distances are annotated per trace. Data represent mean ± SD of biological triplicates (n = 3). (**d**) Correlation between predicted interdomain distance and relative indigoidine production after 48 h. The dashed line indicates a linear regression fit.

To see a possible correlation between protein fragment distance and pigment production, we plot the both parameter against each other. With an R^2^ of 0.7, shorter interdomain distances seem to be associated with higher pigment yields (**Fig. 4d**).

Sequence inversion of one SYNZIP partner disrupts the parallel heptad repeat register, forcing the two partners to reorient into an antiparallel configuration. Simultaneous inversion of both partners restores the original parallel orientation by re-establishing the initial residue register. Interaction specificity in SYNZIPs is critically determined by the register of hydrophobic core ‘a’ and ‘d’ residues and electrostatic contacts at heptad positions g and e (**Supplementary Fig. S2**). When the amino acid sequence of one SYNZIP partner is reversed, the order of residues at these heptad positions is inverted, disrupting the original parallel interaction register. To restore the favorable heptad contact pattern required for stable heterodimerization, the two partners have to re-align in an antiparallel orientation.

A notable exception to this suggested inversion logic is the SZ17:SZ18 pair. A possible explanation is, that in contrast to other SYNZIP pairs, whose specificity is encoded by a specific sequence of electrostatic heptad contacts. In contrast, in the SZ17:SZ18 interaction SZ17 presenting only basic residues and SZ18 presenting only acidic residues at the heptad positions ‘g’ and ‘e’. Consistent with this reasoning, the original FRET analysis by Thompson et al. indicated that SZ17:SZ18 adopts a mixture of parallel and antiparallel orientations in solution^25^, indicating that both geometries are energetically accessible for this SYNZIP pair.

Taken together, the results obtained from our sequence inversion series of SZ1:SZ2 and SZ5:SZ6 consistently demonstrate that inversion of one SYNZIP partner converts parallel pairs into antiparallel configurations, thereby substantially improving performance in the Indi2GO system. These results demonstrate that split-IndC reconstitution efficiency depends on both binding efficiency and domain distance.

### Transferability to other NRPS systems

To assess whether the design principles established in the indigoidine system extend beyond the specific biosynthetic context, all SYNZIP configurations were additionally evaluated in the orthogonal XtpS NRPS system. XtpS is a non-ribosomal peptide synthetase that produces the xenotetrapeptide (XTP, cyclo[vLvV]) through the sequential condensation of four amino acid building blocks^30^. Similar to the indigoidine system, the XtpS enzyme can be split at an XUT^I^ site, with the two resulting fragments unable to reconstitute catalytic activity when separated, thereby abolishing XTP production. Consequently, the production of XTP serves as a quantitative reporter for the efficiency of PPI-mediated complementation in this system.

Quantitative analysis of XTP production was performed by LC-MS/MS analysis of *m/z* = 411.28 [M+H]^+^, which corresponds to the molecular weight of XTP. As a positive control, the covalently linked full-length XtpS was used, while the XUT^I^ split variant without any PPI fusion partner served as a negative control (**Fig. 5**).

**Figure 5.**
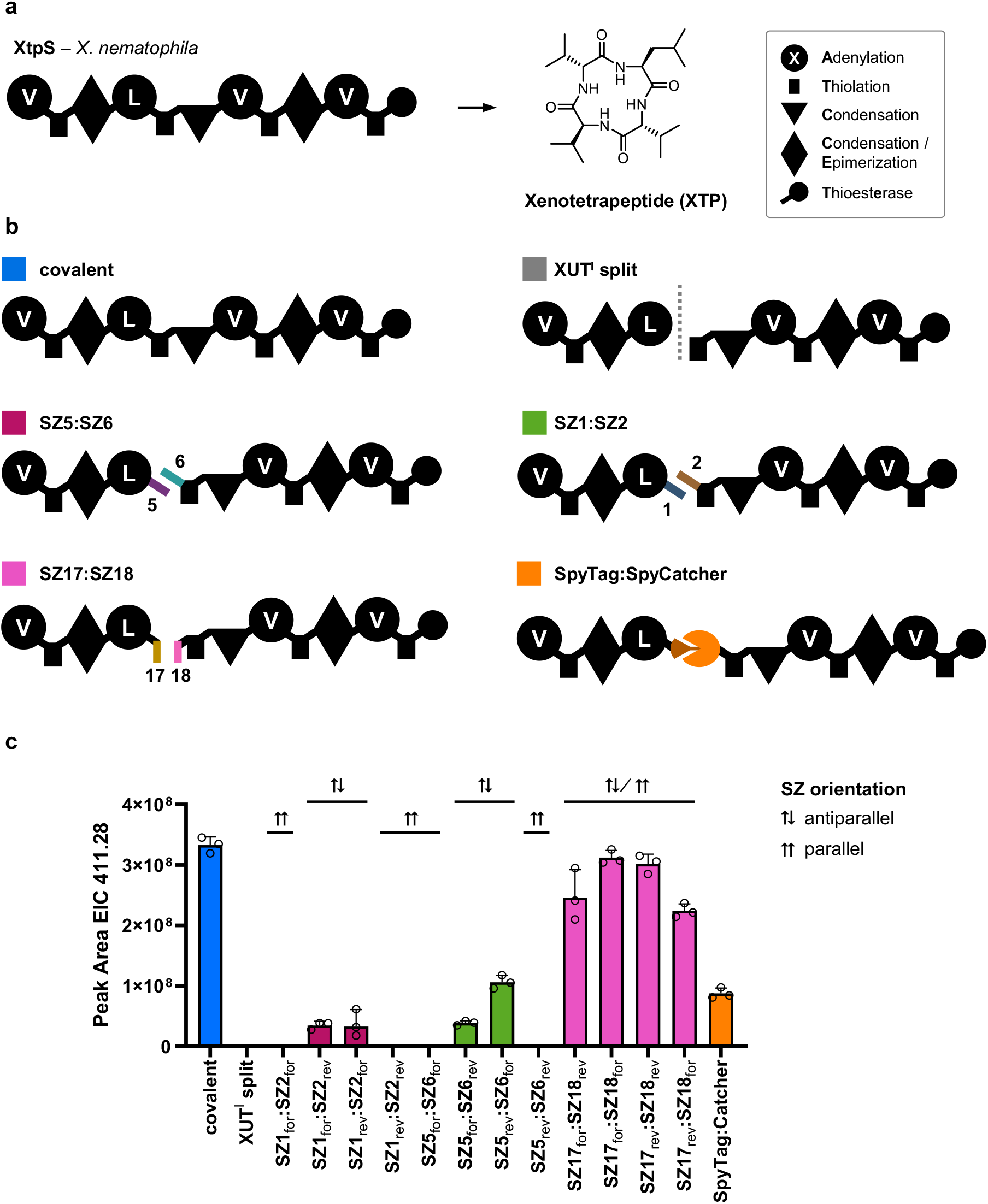
Validation of novel antiparallel SYNZIP configurations in the complex NRPS system XtpS. (**a**) Domain architecture of the xenotetrapeptide synthetase XtpS and the chemical structure of XTP (**b**) Schematic of the split-XtpS construct architecture with all four orientation combinations of SYNZIPS SZ1:SZ2, SZ5:SZ6 and SZ17:SZ18 inserted at an XUT^I^ site, illustrating the parallel (SZ1_for_:SZ2_for_, SZ1_rev_:SZ2_rev_) and antiparallel (SZ1_for_:SZ2_rev_, SZ1_rev_:SZ2_for_) configurations. For SZ17:SZ18, both configurations are possible, with the antiparallel configuration assumed to be the active one. (**c**) Quantification of XTP production by LC-MS peak area analysis of m/z = 411.28 [M+H]+ for all tested SYNZIPS orientation combinations at 48 h post-induction. Data represent mean ± SD of biological triplicates (n = 3).

The results demonstrate that the quantitative correlations observed in the indigoidine system are transferable to other NRPS systems. Parallel configurations of both SYNZIP pairs (SZ1:SZ2 and SZ5:SZ6) failed to restore detectable XTP production, while all antiparallel configurations consistently enabled XTP synthesis. Notably, the antiparallel SZ17:SZ18 pair restored XTP production to levels comparable to the covalent positive control, with the original SZ17_for_:SZ18_for_ configuration exhibiting the highest yield. SZ17:SZ18 can adopt both configurations as described by Thompson *et al*., with the antiparallel configuration assumed to be the active one^25^. The SpyTag:SpyCatcher system also enabled efficient XTP production, achieving yields comparable to those observed with the parallel SZ1:SZ2 pair. These findings confirm that the geometric principles governing split-NRPS reconstitution are not specific to the indigoidine biosynthetic machinery but represent general constraints applicable to diverse NRPS systems.

## CONCLUSION

PPIs are fundamental regulators of cellular function, governing processes from receptor dimerization to signal transduction. The dimerization of receptors such as G-protein coupled receptors (GPCRs) and receptor tyrosine kinases (RTKs) is essential for cellular communication^31^, while dysregulated PPIs contribute to numerous diseases^32^. Consequently, PPIs have emerged as important targets for therapeutic intervention^33^.

Controlling PPIs is also essential for synthetic biology applications to bring two enzymes in close proximity for substrate channeling^34^. Substrate channeling also plays an important role in NRPS biosynthesis in order to transfer the peptide intermediate from one NRPS to another. However in context of NRPS engineering, PPI-mediating tools such as SYNZIPs, split inteins, and the SpyTag:SpyCatcher system were characterized previously using expensive, low-throughput analytical workflows.

Indi2GO now provides a rapid, accessible, and high-throughput compatible split-reporter system for benchmarking PPIs in the context of drug discovery and synthetic biology. Compared to established PPI detection platforms such as yeast two-hybrid assays, FRET, BiFC^11^, or splitFAST^35^, Indi2GO offers great simplicity and accessibility. Analogous to BiFC, which is widely favored because it is easy to implement, Indi2GO requires no specialized instrumentation for qualitative assessment, as the blue chromogenic signal is readily visible by the naked eye. Quantitative analysis requires only a standard spectrophotometer, making this system accessible for laboratories without imaging or detection infrastructure. We acknowledge that the colorimetric readout inherently limits sensitivity and dynamic range relative to fluorescence-based systems. Nevertheless, the non-substrate-dependent nature of the indigoidine synthetase makes Indi2GO a practical platform for comparative analysis of PPI strength.

## ACKNOWLEDGEMENTS

This work was supported by the Max-Planck Society.

## AUTHOR CONTRIBUTIONS

PG, CS and MF designed and conducted the experiments and analyzed the data. LS conducted preliminary work. AP was involved in conceptual discussions. PG, MF and CS created all figures and wrote the manuscript with input from all authors. PG and HBB planned and supervised the project.

## Notes

### Competing Interest Statement

The authors have declared no competing interest.

