## Supplementary material for "Split-Indigoidine synthetase as optical reporter for benchmarking protein-protein interactions": Materials and methods and supplementary information.

### Supplementary Tables

**Supplementary References** ...............................**Fehler! Textmarke nicht definiert.**

### MATERIALS AND METHODS

#### Cultivation of strains

*E. coli* DH10B::*mtaA* cells were cultured in LB medium (10 g/L tryptone, 5 g/L yeast extract, 5 g/L NaCl, pH 7.5). Chloramphenicol (34 µg/mL in ethanol) and kanamycin (50 µg/mL, sterile in dd$H_2O$) were added to the overnight cultures. For cryopreservation, overnight cultures were mixed in a ratio of 1:1 (v/v) with 50 % (v/v) glycerol and stored at -70°C.

#### Polymerase chain reaction (PCR)

DNA fragments and vector backbones were amplified by PCR using Q5 High-Fidelity DNA Polymerase (NEB) and primers containing overhangs. The reaction mixture consisted of 10 µL Q5 Reaction Buffer, 1 µL 10 mM dNTPs, 1 µL forward primer (10 µM), 1 µL reverse primer (10 µM), 1 µL template DNA (~20 ng/µL), 0.5 µL Q5 High-Fidelity DNA Polymerase, 3 µL DMSO, and 32.5 µL nuclease-free water. When plasmids served as templates, the template DNA was digested with 1 µL DpnI followed by incubation at 37°C for 20 min. PCR products were analyzed and purified by 1% agarose gel electrophoresis in TAE buffer containing MIDORI Green Advance (0.2 µL/100 mL, NIPPON Genetics), and DNA fragments were subsequently extracted using the Monarch DNA Gel Extraction Kit (NEB).

#### Gibson cloning of biosynthetic gene clusters

Arabinose inducible $P_{BAD}$ promoters were used as *xtpS* and *indC* were cloned onto pACYC and pCOLA backbone plasmids. All constructs were constructed with Gibson assembly using DNA fragments with 30 bp overlapping homologous overhangs. In the assembly reaction using 20 fmol backbone fragment and 4 molar equivalents of each insert fragment were used. Fragments were assembled with NEBuilder HiFi DNA Assembly Master Mix incubating at 50°C for 1 h. Then 1 µL reaction mix was transformed into chemically competent *E. coli* DH10B cells. Plasmids were confirmed by overnight sequencing (Microsynth) using Sanger sequencing according to the companies protocol. Correct plasmids were isolated from overnight cultures using the

Monarch plasmid miniprep kit (New England Biolabs) using the manufacturers protocol.

#### Preparation of chemically competent cells

*E. coli* DH10B colonies were cultured overnight in 5 mL LB medium at 37°C. The overnight culture was subsequently diluted 1:100 in fresh LB medium and grown at 37°C to an OD600 of 0.5. Cells were chilled on ice and harvested by centrifugation at 2500 rpm and 4°C for 10 min. The supernatant was removed and the pellet was resuspended in buffer 1 containing 15% (v/v) glycerol, 2.94 g/L potassium acetate, 1.47 g/L CaCl2·2H2O, 12.1 g/L RbCl, and 7.2 g/L MnCl2·H2O (pH 5.9). The suspension was incubated on ice for 90 min and centrifuged again at 2500 rpm and 4°C for 10 min. The supernatant was removed and cells were resuspended in buffer 2 consisting of 15% glycerol (v/v), 0.42 g/L MOPS, 0.24 g/L RbCl, and 2.21 g/L CaCl2·2H2O (pH 7.0). Aliquots were then prepared and stored at −70°C.

#### Plasmid transformation

For transformation, competent cells were thawed on ice and mixed with approximately 100 ng plasmid DNA. The cells were incubated on ice for 10 min, heat-shocked at 42°C for 45 s, and returned to ice for an additional 2 min. Subsequently, cells were allowed to recover in SOC medium (10 g/L tryptone, 2.5 g/L yeast extract, 0.25 g/L NaCl, 0.09 g/L KCl, 1.0 g/L MgCl2·6H2O, 1.23 g/L $MgSO_4·7H_2O$, 0.9 g/L glucose) for 1 h at 37°C before plating on LB agar supplemented with the appropriate antibiotics.

#### Indigoidine production for assay

Indigoidine production was assessed in *E. coli* DH10B::*mtaA* cultures grown in XPPM medium. Overnight cultures were diluted 1:100 into 200 µL XPPM medium supplemented with 0.2% (w/v) arabinose and the appropriate antibiotics, in standard greiner 96-well flat-bottom plates. Cultures were incubated and monitored continuously for 48 h post-induction using a Tecan Spark microplate reader operating at 25°C with linear shaking at 810 rpm (2 mm amplitude). Absorbance at 600 nm ($OD_{600}$) and 800 nm ($OD_{800}$) were recorded every 15 min for indigoidine quantification as described below.

### Spectrophotometric quantification of indigoidine production

Indigoidine production was quantified spectrophotometrically as described by Myers *et al.* using a cell density-corrected dual-wavelength absorbance method[1]. The absorption maximum of indigoidine was determined experimentally by performing a full-wavelength scan from 400 to 800 nm in 5 nm increments using a Tecan Spark microplate reader microplate reader. The absorption maximum was identified at 600 nm, which was subsequently used as the indigoidine-sensitive wavelength (ODS) for all quantification experiments (**Fig. S2**). A robust reference wavelength (ODR) of 800 nm was utilized for background correction.This was performed by subtracting the mean absorbance of pure culture medium from all OD values prior to further analysis[1]. The scattering contribution of cellular components was estimated using a correction factor δ, determined from a negative control culture under identical expression conditions. In contrast to previous reports[2] in which uninduced cultures were used to determine δ, the XUT$^{I}$ split-IndC variant lacking any PPI fusion partner was employed as the negative control in this study. This substitution was necessary as the arabinose-inducible $P_{BAD}$ promoter used for construct expression exhibited leaky basal transcription in the absence of inducer, resulting in low but detectable indigoidine production in uninduced full-length IndC cultures. As the XUT$^{I}$ split-IndC variant produced no detectable indigoidine under identical expression conditions (**Fig. 1**), it provided a more accurate estimate of the biomass-only scattering contribution at ODS, and was therefore used as the reference for δ determination in all subsequent experiments:

$$\delta = \frac{\mathrm{OD}_{600\ (\mathrm{reference})}}{\mathrm{OD}_{800\ (\mathrm{reference})}} \tag{1}$$

The biomass-corrected indigoidine absorbance was then calculated as:

$$\mathrm{X} = \mathrm{OD}_{600(\mathrm{sample})} - (\delta \cdot \mathrm{OD}_{800(\mathrm{sample})}) \tag{2}$$

Relative indigoidine production was expressed as a percentage of the full-length IndC positive control:

$$\text{Relative indigoidine production (\%)} = \frac{X_{\text{sample}}}{X_{\text{positive control}}} \cdot 100\% \quad (3)$$

All absorbance measurements were performed using a Tecan Spark microplate reader microplate reader in standard 96-well flat-bottom plates. Indigoidine production was monitored continuously over 48 h post-induction, with absorbance measurements recorded at 15 min intervals at both 600 nm and 800 nm. Unless otherwise stated, the 48 h timepoint was used for endpoint quantification and comparison between constructs.

**XTP expression and extraction**

100 µL of overnight culture was inoculated into 10 mL XPPM medium (10 g/L glycerol, 20 mL/L M9 salt A, 20 mL/L M9 salt B, 20 g/L; M9 salt A: 350 g/L $K_2HPO_4$, 100 g/L $KH_2PO_4$; M9 salt B: 5 g/L $MgSO_4$, 50 g/L $(NH_4)_2SO_4$, 29.4 g/L sodium citrate) supplemented with 2 mL/L vitamins (6 mg/L biotin, 10 mg/L folic acid, 2.3 g/L nicotinic acid, 1.2 g/L panthothenic acid, 200 mg/L p-aminobenzoic acid, 12 g/L pyridoxine-HCl, 100 mg/L riboflavin, 1 g/L Thiamine-HCl, 20 mg/L vitamin B12) and 1 mL/L trace elements (10 mg/L $(NH_4)_6Mo_7O_{24}\cdot 4H_2O$, 10 mg/L $CuCl_2\cdot 2H_2O$, 10 mg/L $MnCl_2\cdot 4H_2O$, 10 mg/L $Na_2B_4O_7\cdot 10H_2O$, 200 mg/L $FeCl_3\cdot 6H_2O$, 40 mg/L $ZnCl_2$). Arabinose and antibiotics were used as described above. Cultures were grown at 25°C for 48 h shaken at 200 rpm in 10 mL flasks. Cultures were diluted 1:5 in MeOH:ACN (1:1). 1 mL of the extract was centrifuged at 13000 rpm for 20 minutes and 100 µL of the supernatant was measured on the HPLC-MS.

**LC-MS/MS measurement**

HPLC-MS analysis was conducted using an Agilent 1290 Infinity II UPLC system equipped with a Waters ACQUITY Premier BEH C18 column (186009453) coupled to a Bruker AmaZon Speed ESI-IT mass spectrometer. Samples (5 µL injection volume) were separated via a 16 min gradient from 5% to 95% acetonitrile (+0.1% formic acid) in water (+0.1% formic acid). Mass spectra were acquired in positive ionization mode (m/z 100–1200, cone voltage 4500 V), and data processing was performed with Bruker's DataAnalysis software.

For relative quantification of xenotetrapeptide production, biological triplicates were analyzed. Peak areas were calculated from the extracted ion chromatogram (EIC) of Xenotetrapeptide (m+1 = 411.28) and visualized in GraphPad Prism 11.

### SUPPLEMENTARY TABLES

#### Table S1. | *E. coli* strains used in this work.

| Strain | Genotype | Reference |
|---|---|---|
| *E. coli* DH10B | F-mcrA, Δ(mrr-hsdRMS-mcrBC), Φ80lacZΔM15, ΔlacX74, recA1, endA1, araD139, Δ(ara leu)7697, galU, galK, rpsL, nupG, λ–, entD::mtaA | [3] |
| *E. coli* DH10B::mtaA | DH10B with mtaA from pCK_mtaA Δ entD | [4] |

#### Table S2. | *Photorhabdus and Xenorhabdus* strains with respective BGC and natural products used in this work.

| Strain | | Gene | Locus tag | GenBank | NP, PubChem CID |
|---|---|---|---|---|---|
| *Xenorhabdus nematophila* | ATCC 19061 | *xtpS* | XNC1_2022 | FN667742 | Xenotetrapeptide 145720651 |
| *Photorhabdus laumondii* TT01 | DSM 15139 | *indC* | PluTT01m_11290 | CP024901.1 | Indigoidine, 193349 |

#### Table S3. | Natural products used this work.

| Name | PubChem CID | SMILES | Exact mass (Da) |
|---|---|---|---|
| Xenotetrapeptide (XTP) | 145720651 | CC(C)C[C@H]1C(=O)N[C@@H](C(=O)N[C@H](C(=O)N[C@@H](C(=O)N1)C(C)C)C(C)C)C(C)C | 410.28 |
| Indigoidine | 193349 | C1=C(C(=O)NC(=O)C1=N)C2=C(NC(=O)C(=C2)N)O | 248.05 |

**Table S4. | Plasmids cloned in this work.**

| Plasmid | Fragment name | Forward primer | Reverse primer | Template | Size (bp) | Resis-tance |
|---|---|---|---|---|---|---|
| pMF1 | pcrMF001 | oMF1F | oMF2R | pLS70 (indC) | 2883 | Cm |
| | pcrMF002 | oMF3F | oMF4R | pCS90-1 | 5349 | |
| pMF2 | pcrMF003 | oMF5F | oMF6R | pLS70 (indC) | 1086 | Kan |
| | pcrMF004 | oMF7F | oMF8R | pCS91 | 3435 | |
| pMF3 | pcrMF005 | oMF9F | oMF6R | pLS70 (indC) | 1086 | Kan |
| | pcrMF006 | oMF7F | oMF10R | pJW76 | 3435 | |
| pMF4 | pcrMF007 | oMF1F | oMF11R | pLS70 (indC) | 2883 | Cm |
| | pcrMF008 | oMF12F | oMF4R | pCS90 | 5223 | |
| pMF5 | pcrMF009 | oMF13F | oMF6R | pLS70 (indC) | 1086 | Kan |
| | pcrMF010 | oMF7F | oMF14R | pCS91 | 3312 | |
| pMF6 | pcrMF011 | oMF1F | oMF15R | pLS70 (indC) | 3120 | Cm |
| | pcrMF008 | oMF12F | oMF4R | pCS90 | 5223 | |
| pMF7 | pcrMF012 | oMF16F | oMF6R | pLS70 (indC) | 849 | Kan |
| | pcrMF010 | oMF7F | oMF14R | pCS91 | 3312 | |
| pMF8 | pcrMF13 | oMF1F | oMF17R | pLS70 (indC) | 2883 | Cm |
| | pcrMF14 | oMF18F | oMF4R | pIN05 | 5547 | |
| pMF9 | pcrMF15 | oMF19F | oMF6R | pLS70 (indC) | 1129 | Kan |
| | pcrMF16 | oMF7F | oMF20R | pIN06 | 3385 | |
| pMF13 | pcrMF20 | oMF1F | oMF24R | pLS70 (indC) | 2958 | Cm |
| | pcrMF08 | oMF12F | oMF4R | pCS90 | 5223 | |
| pMF14 | pcrMF21 | oMF25F | oMF6R | pLS70 (indC) | 1011 | Kan |
| | pcrMF010 | oMF7F | oMF14R | pCS91 | 3312 | |
| pMF17 | pcrMF24 | oMF28F | oMF29R | pMF1 | 8190 | Cm |
| pMF18 | pcrMF25 | oMF30F | oMF31R | pMF2 | 4497 | Kan |
| pMF19 | pcrMF26 | oMF32F | oMF33R | pMF3 | 4479 | Kan |
| pMF20 | pcrMF27 | oMF34F | oMF35R | pMF1 | 8199 | Cm |
| pMF21 | pcrMF28 | oMF36F | oMF37R | pMF2 | 8199 | Kan |
| pMF22 | pcrMF29 | oMF38F | oMF39R | pMF3 | 4488 | Kan |
| pMF23 | pcrMF30 | oMF40F | oMF41R | pMF1 | 8208 | Cm |
| pMF24 | pcrMF31 | oMF42F | oMF43R | pMF2 | 4497 | Kan |
| pMF25 | pcrMF32 | oMF44F | oMF45R | pMF3 | 4497 | Kan |
| pMF26 | pcrMF33 | oMF47F | oMF46R | pMF17 | 8151 | Cm |
| pMF27 | pcrMF34 | oMF51F | oMF50R | pMF18 | 4452 | Kan |
| pMF28 | pcrMF35 | oMF53F | oMF52R | pMF19 | 4470 | Kan |
| pMF29 | pcrMF36 | oMF47F | oMF48R | pMF17 | 8130 | Cm |
| pMF30 | pcrMF37 | oMF54F | oMF50R | pMF18 | 4419 | Kan |
| pMF31 | pcrMF38 | oMF55F | oMF52R | pMF19 | 4419 | Kan |

| | | | | | | |
|---|---|---|---|---|---|---|
| pMF32 | pcrMF39 | oMF47F | oMF49R | pMF17 | 8190 | Cm |
| pMF33 | pcrMF40 | oMF56F | oMF50R | pMF18 | 4389 | Kan |
| pMF34 | pcRMF41 | oMF57F | oMF52R | pMF19 | 4389 | Kan |
| pMF35 | pcrMF42 | oMF58F | oMF59R | pCS098 | 204 | Cm |
| | pcrMF43 | oMF60F | oMF61R | pMF17 | 8055 | |
| pMF36 | pcrMF44 | oMF62F | oMF63R | pCS099 | 210 | Kan |
| | pcrMF45 | oMF64F | oMF65R | pMF18 | 4344 | |
| pMF37 | pcrMF46 | oMF66F | oMF67R | pCS100 | 210 | Cm |
| | pcrMF43 | oMF60F | oMF61R | pMF17 | 8055 | |
| pMF38 | pcrMF47_B | oMF68F_B | oMF69R_B | pCS101 | 215 | Kan |
| | pcrMF45 | oMF64F | oMF65R | pMF18 | 4344 | |
| pMF39 | pcrMF68 | oMF84F | oCS254R | pMF17 | 2268 | Cm |
| | pcrMF69 | oCS253F | oMF85R | pMF17 | 5981 | |
| pMF40 | pcrMF49 | oMF72F | oMF73R | pCS103 | 224 | Kan |
| | pcrMF45 | oMF64F | oMF65R | pMF18 | 4344 | |
| pMF41 | pcrMF50 | oMF74F | oMF75R | pCS104 | 195 | Cm |
| | pcrMF43 | oMF60F | oMF61R | pMF17 | 8055 | |
| pMF42 | pcrMF51 | oMF76F | oMF77R | pCS105 | 225 | Kan |
| | pcrMF45 | oMF64F | oMF65R | pMF18 | 4344 | |
| pMF52 | pcrMF65 | oMF103F | oMF104R | pMF23 (SpyTag (Klebe et. Al.) | 8151 | Cm |
| pMF53 | pcrMF66 | oMF105F | oMF106R | pMF24 | 4365 | Kan |
| | fMF001 | | | | 399 | |
| pMF64 | pcrMF73 | oMF119F | oMF120R | pXtpS | 4896 | Cm |
| | pcrMF74 | oMF121F | oMF122R | pMF17 | 5418 | |
| pMF65 | pcrMF75a | oMF123F | oIN135R | pXtpS | 3249 | Kan |
| | pcrMF75b | oIN136F | oMF124R | pXtpS | 4296 | |
| | pcrMF76 | oMF125F | oMF126R | pMF19 | 3504 | |
| pMF66 | pcrMF73 | | | | 4896 | Cm |
| | pcrMF78 | oMF127F | oMF128R | pMF59 | 5418 | |
| pMF67 | pcrMF75a | | | | 3249 | Kan |
| | pcrMF75b | | | | 4296 | |
| | pcrMF79 | oMF129F | oMF130R | pMF18 | 3504 | |
| pMF68 | pcrMF73 | | | | 4896 | Cm |
| | pcrMF80 | oMF131F | oMF132R | pMF35 | 5433 | |
| pMF69 | pcrMF75a | | | | 3249 | Kan |
| | pcrMF75b | | | | 4296 | |
| | pcrMF81 | oMF133F | oMF134R | pMF36 | 3531 | |
| pMF70 | pcrMF73 | | | | 4896 | Cm |
| | pcrMF82 | oMF135F | oMF136R | pMF37 | 5433 | |
| pMF72 | pcrMF73 | | | | 4896 | Cm |

| | | | | | | |
|---|---|---|---|---|---|---|
| | pcrMF85 | oMF140F | oMF141R | pMF39 | 5424 | |
| pMF73 | pcrMF75a | | | | 3249 | Kan |
| | pcrMF75b | | | | 4296 | |
| | pcrMF86 | oMF143F | oMF144R | pMF40 | 3543 | |
| pMF74 | pcrMF73 | | | | 4896 | Cm |
| | pcrMF87 | oMF145F | oMF146R | pMF41 | 5424 | |
| pMF75 | pcrMF75a | | | | 3249 | Kan |
| | pcrMF75b | | | | 4296 | |
| | pcrMF88 | oMF147F | oMF148R | pMF42 | 3543 | |
| pMF76 | pcrMF90 | oMF150F | oMF151R | pXtpS | 4896 | Cm |
| | pcrMF91 | oMF152F | oMF153R | pMF52 | 5358 | |
| pMF77 | pcrMF92 | oMF154F | oIN135R | pXtpS | 3249 | Kan |
| | pcrMF93 | oIN136F | oMF155R | pXtpS | 4296 | |
| | pcrMF94 | oMF156F | oMF157R | pMF53 | 3738 | |

**Table S5. | Oligonucleotides used in this work.**

| Name | Sequence 5' -> 3' | Length |
|---|---|---|
| oMF1 | TTTTTTTGGGCTAACAGGAGGAATTCCATGTTAGAAAATAATATTACACAATGTGACTCAATCAATGATGTTTATCTTAA | 80 |
| oMF2 | CTTTTTCGATTTTAATTCCTCCTTCTCGTTTAGACGCTGTGTTGAAACATTCTCCACG | 58 |
| oMF3 | AACGAGAAGGAGGAATTAAAATCGAAAAAGGCTG | 34 |
| oMF4 | CATGGAATTCCTCCTGTTAGCCCAAAAAAACG | 32 |
| oMF5 | GCGAAACTGGAGCGTGAAGAAGCGTACTTCTTGGTGCCATTACATACAGATACTGAAATAAGGC | 64 |
| oMF6 | CACATTATACGAGCCGATGATTAATTGTCAGATTATTTTCTCAATCTCAGCAACACCTTCACTTTC | 66 |
| oMF7 | TGACAATTAATCATCGGCTCGTATAATGTGTGGA | 34 |
| oMF8 | GAAGTACGCTTCTTCACGCTCCAGTTTC | 28 |
| oMF11 | CACATTATACGAGCCGATGATTAATTGTCATAGACGCTGTGTTGAAACATTCTCCACG | 58 |
| oMF12 | TGACAATTAATCATCGGCTCGTATAATGTGTGGAATTG | 38 |
| oMF13 | TTTTTTTGGGCTAACAGGAGGAATTCCATGTTGGTGCCATTACATACAGATACTGAAATAAGGCTTGG | 68 |
| oMF14 | CATGGAATTCCTCCTGTTAGCCCAAAAAAACG | 32 |
| oMF15 | CACATTATACGAGCCGATGATTAATTGTCAAGAGTCTGTCTGTTCAATCCACTTAGCCAATTC | 63 |

| | | |
|---|---|---|
| oMF16 | TTTTTTTGGGCTAACAGGAGGAATTCCATGAAAACAATATCAAGATTAATTTTATTGAATCAGGCAAGCAAAGACCCC | 78 |
| oMF17 | TTCGCTGCTACCAACCAGACAACAAGGGTTTAGACGCTGTGTTGAAACATTCTCCACG | 58 |
| oMF18 | AACCCTTGTTGTCTGGTTGGTAGCAGC | 27 |
| oMF19 | GGATACCTATGATATTCAGACCAGCACCCATAACTTTTTCGCCAATGATATTCTGGTGCATAATAGCGAAATTTTGGTGCCATTACATACAGATACTGAAATAAGGCTTGG | 111 |
| oMF20 | TCTGAATATCATAGGTATCCTCGTTGCTCACAATTTTACGGCTTTTCAGTTTGCCGATATAGGTTTTGGCTTCCATGGAATTCCTCCTGTTAGCCCAAAAAAACG | 105 |
| oMF23 | CACATTATACGAGCCGATGATTAATTGTCATACTGAATCCCATTTCAGTACTTCCATCCAAATTTTTCC | 69 |
| oMF24 | TTTTTTTGGGCTAACAGGAGGAATTCCATGTCTGCCCTCGATGATTTTTTCGAAAGTGG | 59 |
| oMF25 | TTTTTTTGGGCTAACAGGAGGAATTCCATGTCTGCCCTCGATGATTTTTTCGAAAGTGG | 59 |
| oMF28 | GGCAGTGGTAACGAGAAGGAGGAATTAAAATCGAAAAAGGCTG | 43 |
| oMF29 | ACCACTGCCTAGACGCTGTGTTGAAACATTCTCCACG | 37 |
| oMF30 | GGCAGTGGTTTGGTGCCATTACATACAGATACTGAAATAAGGC | 43 |
| oMF31 | ACCACTGCCGAAGTACGCTTCTTCACGCTCCAGTTTC | 37 |
| oMF32 | GGCAGTGGTTTGGTGCCATTACATACAGATACTGAAATAAGGC | 43 |
| oMF33 | ACCACTGCCTGAGATAGCTGCAGTCAGCTCGTTATCAAGG | 40 |
| oMF34 | GTGGTGGGAGCGGTAACGAGAAGGAGGAATTAAAATCGAAAAAGGCTG | 48 |
| oMF35 | TCCCACCACTGCCTAGACGCTGTGTTGAAACATTCTCCACG | 41 |
| oMF36 | GTGGTGGGAGCGGTTTGGTGCCATTACATACAGATACTGAAATAAGGC | 48 |
| oMF37 | TCCCACCACTGCCGAAGTACGCTTCTTCACGCTCCAGTTTC | 41 |
| oMF38 | GTGGTGGGAGCGGTTTGGTGCCATTACATACAGATACTGAAATAAGGC | 48 |
| oMF39 | TCCCACCACTGCCTGAGATAGCTGCAGTCAGCTCGTTATCAAGG | 44 |
| oMF40 | GGGAGCGGTGGTTCTGGCAACGAGAAGGAGGAATTAAAATCGAAAAAGGCTG | 52 |
| oMF41 | ACCGCTCCCACCACTGCCTAGACGCTGTGTTGAAACATTCTCCACG | 46 |
| oMF42 | GGGAGCGGTGGTTCTGGCTTGGTGCCATTACATACAGATACTGAAATAAGGC | 52 |

| | | |
|---|---|---|
| oMF43 | ACCGCTCCCACCACTGCCGAAGTACGCTTCTTCACGCTCCAGTTTC | 46 |
| oMF44 | GGGAGCGGTGGTTCTGGCTTGGTGCCATTACATACAGATACTGAAATAAGGC | 52 |
| oMF45 | ACCGCTCCCACCACTGCCTGAGATAGCTGCAGTCAGCTCGTTATCAAGG | 49 |
| oMF46 | TAATTGTCAGGCGATCTTTTGCTTTAATTGTTCACGCTTC | 40 |
| oMF47 | TGACAATTAATCATCGGCTCGTATAATGTGTGGAATTGTGAGC | 43 |
| oMF48 | TAATTGTCACTGTTTTAACTGTTCGATGCGATTACGCAATTC | 42 |
| oMF49 | TAATTGTCATTCAGCCTTTTTCGATTTTAATTCCTCCTTCTCG | 43 |
| oMF50 | CATGGAATTCCTCCTGTTAGCCCAAAAAAACGGG | 34 |
| oMF51 | AATTCCATGCTGGCGCGTCTGGAAAACGAAAACG | 34 |
| oMF52 | CATGGAATTCCTCCTGTTAGCCCAAAAAAAC | 31 |
| oMF53 | AATTCCATGGCCTTGGACCGCGAGTTAAATGCC | 33 |
| oMF54 | AATTCCATGCTCGAAAAAGACATCGCGAACCTGGAAC | 37 |
| oMF55 | AATTCCATGAAAGAGCTGCGTGCCAACGAAAACGAAC | 37 |
| oMF56 | AATTCCATGGACCTGGCGAAACTGGAGCGTGAAG | 34 |
| oMF57 | AATTCCATGCGCGCCCTTGATAACGAGCTGAC | 32 |
| oMF58 | AATGTTTCAACACAGCGTCTAGGCAGTGGTAACCTGGTTGCGCAGCTCGAAAACG | 55 |
| oMF59 | CCACACATTATACGAGCCGATGATTAATTGTCATTCTTCGATTTTCTTACGCAGATTCGCGATTTC | 66 |
| oMF60 | TGACAATTAATCATCGGCTCGTATAATGTGTGG | 33 |
| oMF61 | ACCACTGCCTAGACGCTGTGTTGAAACA | 28 |
| oMF62 | TTTTTTTGGGCTAACAGGAGGAATTCCATGGCGCGTAACGCGTATCTGCGTAAG | 54 |
| oMF63 | ATCTGTATGTAATGGCACCAAACCACTGCCCTGTTCGTGAGACGCAACTTCGTTTTCG | 58 |
| oMF64 | GGCAGTGGTTTGGTGCCATTACATACAGATAC | 32 |
| oMF65 | CATGGAATTCCTCCTGTTAGCCCAAAAAAACGGGTA | 36 |
| oMF66 | AATGTTTCAACACAGCGTCTAGGCAGTGGTGAAGAAATCAAAAAGCGTCTGAATGCGATCG | 61 |
| oMF67 | CCACACATTATACGAGCCGATGATTAATTGTCAGTTCAGAACCGCCTGGAGTTCGTTTTC | 60 |
| oMF68 | ATCTGTATGTAATGGCACCAAACCACTGCCCAGGAACACTCTGCGGTTGAAAACGAACTC | 60 |
| oMF69 | TTTTTTTGGGCTAACAGGAGGAATTCCATGCGCACGGTTCGCATACAGACGCTTTTTG | 58 |
| oMF70 | AATGTTTCAACACAGCGTCTAGGCAGTGGTAACACCGTTAAAGAACTGAAAAACTACATCCAGG | 64 |
| oMF71 | CACATTATACGAGCCGATGATTAATTGTCACTCGAATTTGTGAGCCGCCAGTTCG | 55 |

| | | |
|---|---|---|
| oMF72 | TTTTTTTGGGCTAACAGGAGGAATTCCATGCAAAAAGT TGCGCAGCTGAAAAACCGTGTTG | 61 |
| oMF73 | ATCTGTATGTAATGGCACCAAACCACTGCCACGCGCAA CGTCACGTTCCAGATTTG | 56 |
| oMF74 | AATGTTTCAACACAGCGTCTAGGCAGTGGTGAGTTCAA ACACGCTGCGCTGGAATTC | 57 |
| oMF75 | CACATTATACGAGCCGATGATTAATTGTCAGTTGGTAA CTTTTTCCAGTTTGTTGTAGATCTGC | 64 |
| oMF76 | TTTTTTTGGGCTAACAGGAGGAATTCCATGCGTGCGGT TGACCGTGAACTGAATG | 55 |
| oMF77 | ATCTGTATGTAATGGCACCAAACCACTGCCTTGTTTAA CCGCCTGCAGTTTGTTACGAAC | 60 |
| oMF103 | ATCGTGATGGTGGACGCCTACAAGCGTTACAAGTGAC AATTAATCATCGGCTCGTATAATGTGTGG | 66 |
| oMF104 | GTCCACCATCACGATATGAGGCACGCCACGGCCAGAA CCACCGCTCCCACC | 51 |
| oMF105 | GGCAGTGGTGGGAGCGGTGGTTC | 23 |
| oMF106 | CATGGAATTCCTCCTGTTAGCCCAAAAAAACG | 32 |
| oMF119F | AAAGATAGCATGGCTAAAAAGGGAATTATCTTTGACGC | 38 |
| oMF120R | AATCTGGCGGGCGAAAGCCTCTTC | 24 |
| oMF121F | CCGGGGGAAGAGGCTTTCGCCCGCCAGATTGGCAGT GGTAACGAGAAGGAGGAATTAAAATCG | 63 |
| oMF122R | GATAATTCCCTTTTTAGCCATGCTATCTTTCATGGAATT CCTCCTGTTAGCCCAAAAAAACG | 62 |
| oMF123F | TATGTTGCGCCACAAGGAGAAATGGAAATCG | 31 |
| oMF124R | CAGCGCCTCCACTTCGCAATTCATTGC | 27 |
| oMF125F | TTGGCAATGAATTGCGAAGTGGAGGCGCTGTGACAATT AATCATCGGCTCGTATAATGTGTGGA | 64 |
| oMF126R | GATTTCCATTTCTCCTTGTGGCGCAACATAACCACTGC CTGAGATAGCTGCAGTCAG | 57 |
| oMF127F | CCGGGGGAAGAGGCTTTCGCCCGCCAGATTGGCAGT GGTAAATATGCAGAGATAGAGAAACG | 62 |
| oMF128R | GATAATTCCCTTTTTAGCCATGCTATCTTTCATGGAATT CCTCCTGTTAGCCCAAAAAAACG | 62 |
| oMF129F | TTGGCAATGAATTGCGAAGTGGAGGCGCTGTGACAATT AATCATCGGCTCGTATAATGTGTGG | 63 |
| oMF130R | GATTTCCATTTCTCCTTGTGGCGCAACATAACCACTGC CGAAGTACGCTTCTTCAC | 56 |
| oMF131F | CCGGGGGAAGAGGCTTTCGCCCGCCAGATTGGCAGT GGTAACCTGGTTGCGC | 52 |
| oMF132R | GATAATTCCCTTTTTAGCCATGCTATCTTTCATGGAATT CCTCCTGTTAGCCCAAAAAAACG | 62 |
| oMF133F | TTGGCAATGAATTGCGAAGTGGAGGCGCTGTGACAATT AATCATCGGCTCGTATAATGTGTGG | 63 |
| oMF134R | GATTTCCATTTCTCCTTGTGGCGCAACATAACCACTGC CCTGTTCGTGAGACG | 53 |

| | | |
|---|---|---|
| oMF135F | CCGGGGGAAGAGGCTTTCGCCCGCCAGATTGGCAGT GGTGAAGAAATCAAAAAGCGTC | 58 |
| oMF136R | AGATAATTCCCTTTTTAGCCATGCTATCTTTCATGGAAT TCCTCCTGTTAGCCCAAAAAAACG | 63 |
| oMF137F | TGAACCAGGAAGATCGTGAACTGCAGCTGAACGACAA AAAACTGCGTGCAATCAAAAAGCGTCTGTATGCGAACC GTGCGGGCAGTGGTTATGTTGCGCCACAAGGAGAAAT GGAAATCG | 120 |
| oMF138F | TTGGCAATGAATTGCGAAGTGGAGGCGCTGTGACAATT AATCATCGGCTCGTATAATGTGTGGAATTGTG | 70 |
| oMF139R | CTTCCTGGTTCAGTTCTTTGATGATCGCGTTCAGACGG TCTTCGATCGCACGGAGTTCGTTTTCAACCGCAGAGTG TTCCTGCATGGAATTCCTCCTGTTAGCCCAAAAAAACG | 114 |
| oMF140F | CCGGGGGAAGAGGCTTTCGCCCGCCAGATTGGCAGT GGTAACACCGTTAAAGAACTG | 57 |
| oMF141R | GATAATTCCCTTTTTAGCCATGCTATCTTTCATGGAATT CCTCCTGTTAGCCCAAAAAAACG | 62 |
| oMF143F | TTGGCAATGAATTGCGAAGTGGAGGCGCTGTGACAATT AATCATCGGCTCGTATAATGTGTGG | 63 |
| oMF144R | GATTTCCATTTCTCCTTGTGGCGCAACATAACCACTGC CACGCGCAACGTCACG | 54 |
| oMF145F | CCGGGGGAAGAGGCTTTCGCCCGCCAGATTGGCAGT GGTGAGTTCAAACACGCTGCGC | 58 |
| oMF146R | GATAATTCCCTTTTTAGCCATGCTATCTTTCATGGAATT CCTCCTGTTAGCCCAAAAAAACG | 62 |
| oMF147F | TTGGCAATGAATTGCGAAGTGGAGGCGCTGTGACAATT AATCATCGGCTCGTATAATGTGTGG | 63 |
| oMF148R | GATTTCCATTTCTCCTTGTGGCGCAACATAACCACTGC CTTGTTTAACCGCCTGC | 55 |
| oMF150F | AAAGATAGCATGGCTAAAAAGGGAATTATCTTTGACG | 37 |
| oMF151R | AATCTGGCGGGCGAAAGCCTCTTCC | 25 |
| oMF152F | CCGGGGGAAGAGGCTTTCGCCCGCCAGATTGGCAGT GGTGGGAGCGGTGGTTCTG | 55 |
| oMF153R | GATAATTCCCTTTTTAGCCATGCTATCTTTCATGGAATT CCTCCTGTTAGCCCAAAAAAACG | 62 |
| oMF154F | TATGTTGCGCCACAAGGAGAAATGGAAATCG | 31 |
| oMF155R | CAGCGCCTCCACTTCGCAATTCATTGC | 27 |
| oMF156F | TTGGCAATGAATTGCGAAGTGGAGGCGCTGTGACAATT AATCATCGGCTCGTATAATGTGTGGA | 64 |
| oMF157R | GATTTCCATTTCTCCTTGTGGCGCAACATAGCCAGAAC CACCGCTCCCACCACTG | 55 |

**Table S6. | Synthetic DNA fragments used in this work.**

| **Fragment Name** | **Sequence 5' -> 3'** |
|---|---|
| fMF001 - SpyCatcher 003 | TTTTTTTGGGCTAACAGGAGGAATTCCATGGTAACCACCTTATCAGGTTTATCAGGTGAGCAAGGTCCGTCCGGTGATATGACAACTGAAGAAGATAGTGCTACCCATATTAAATTCTCAAAACGTGATGAGGACGGCCGTGAGTTAGCTGGTGCAACTATGGAGTTGCGTGATTCATCTGGTAAAACTATTAGTACATGGATTTCAGATGGACATGTGAAGGATTTCTACCTGTATCCAGGAAAATATACATTTGTCGAAACCGCAGCACCAGACGGTTATGAGGTAGCAACTCCAATTGAATTTACAGTTAATGAGGACGGTCAGGTTACTGTAGATGGTGAAGCAACTGAAGGTGACGCTCATACTGGCAGTGGTGGGAGCGGTGGTTCTGGCTTG |

### SUPPLEMENTARY Figures

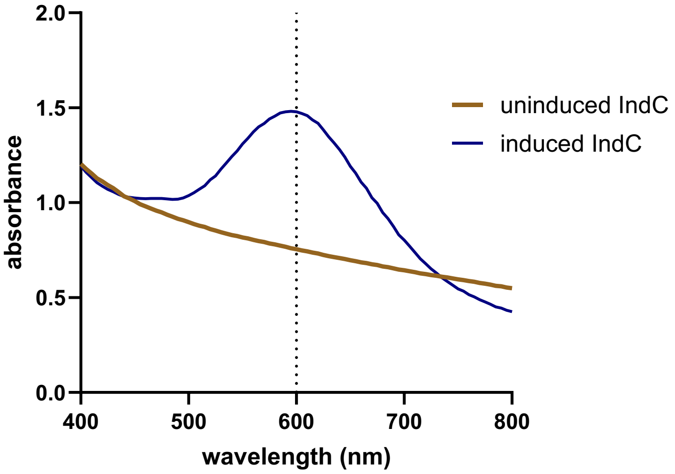

**Figure S1 | Indigoidine absorbance spectrum and biomass contribution analysis.** Absorbance spectrum of indigoidine in bacterial culture measured from 400 to 800 nm in 5 nm increments at 24 h post-induction in XPPM medium. The spectrum shows a pronounced absorption maximum at 600 nm, consistent with the wavelength used for indigoidine quantification. Linear relationship between $OD_{600}$ and $OD_{800}$ in uninduced bacterial cultures used for biomass-corrected absorbance calculation.

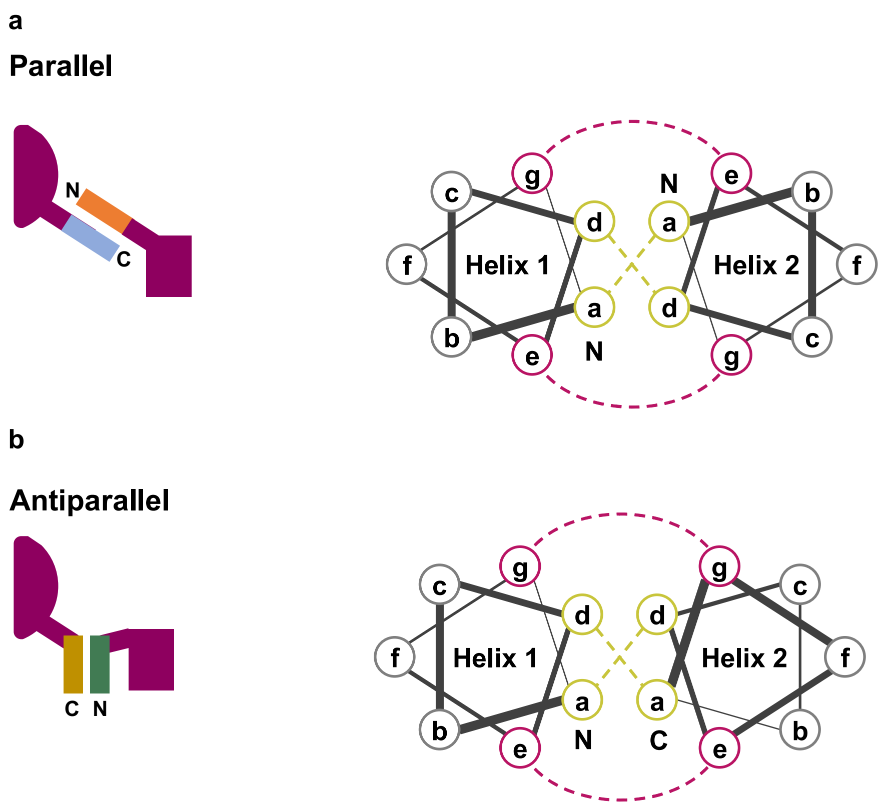

**Figure S2 | Parallel and antiparallel zipper interaction geometry.** (**a**)Schematic representation of parallel SYNZIP interaction showing the heptad repeat pattern (abcdefg)$_n$ with hydrophobic residues at positions a and d and electrostatic residues at positions e and g. In parallel orientation both N-termini and both C-termini of the SYNZIPs interact, placing the fused NRPS fragments at maximal distance from each other. The parallel g'-e' heptad interaction stabilizes the SYNZIP dimer. (**b**) Schematic of antiparallel SYNZIP interaction the N-terminus of one SYNZIP partner interacts with the C-terminus of the other partner, bringing the fused NRPS fragments into close proximity. In antiparallel orientation g'-g' and e'-e' interaction, creating a complementary electrostatic interaction pattern.

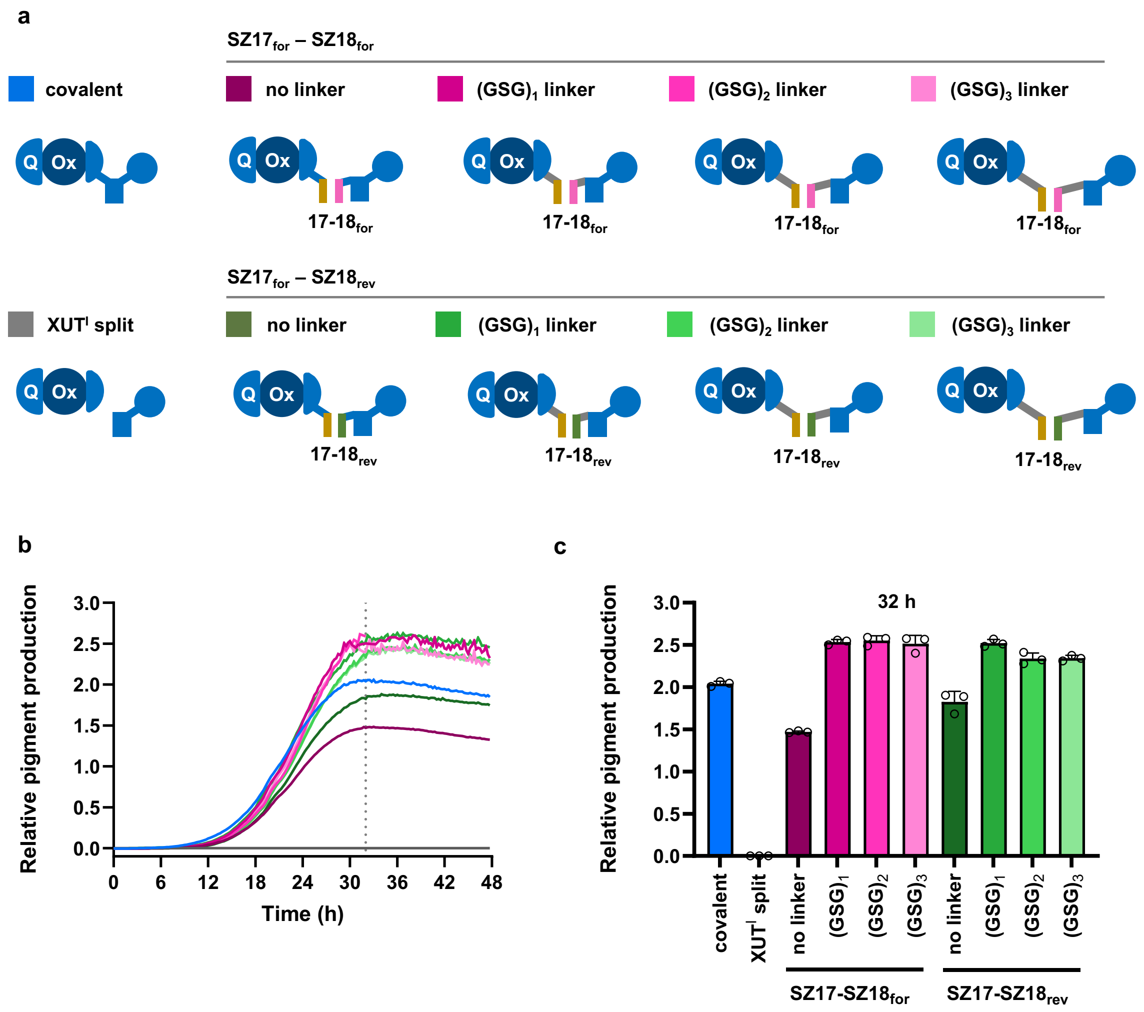

**Figure S3| Linker optimization for SYNZIPs.** (**a**) Schematic of the split-IndC construct with SYNZIP pairs SZ17:SZ18for and SZ17:SZ18rev and GSG linkers of varying length $(GSG)_x$ (x=1, 2, 3) at the $XUT^I$ site. (**b**) Time-course of relative indigoidine production of split-IndC constructs carrying $SZ17:SZ18_{for}$ and $SZ17:SZ18_{rev}$ with increasing GSG linker lengths over 48 h post-induction, normalized to withon GS linker IndC (100%). (**c**) Endpoint quantification of relative indigoidine production at 32 h mean ± SD of biological triplicates (n = 3) for all tested linker configurations. Introduction of a single GSG linker unit substantially increased indigoidine production for both SYNZIPs.

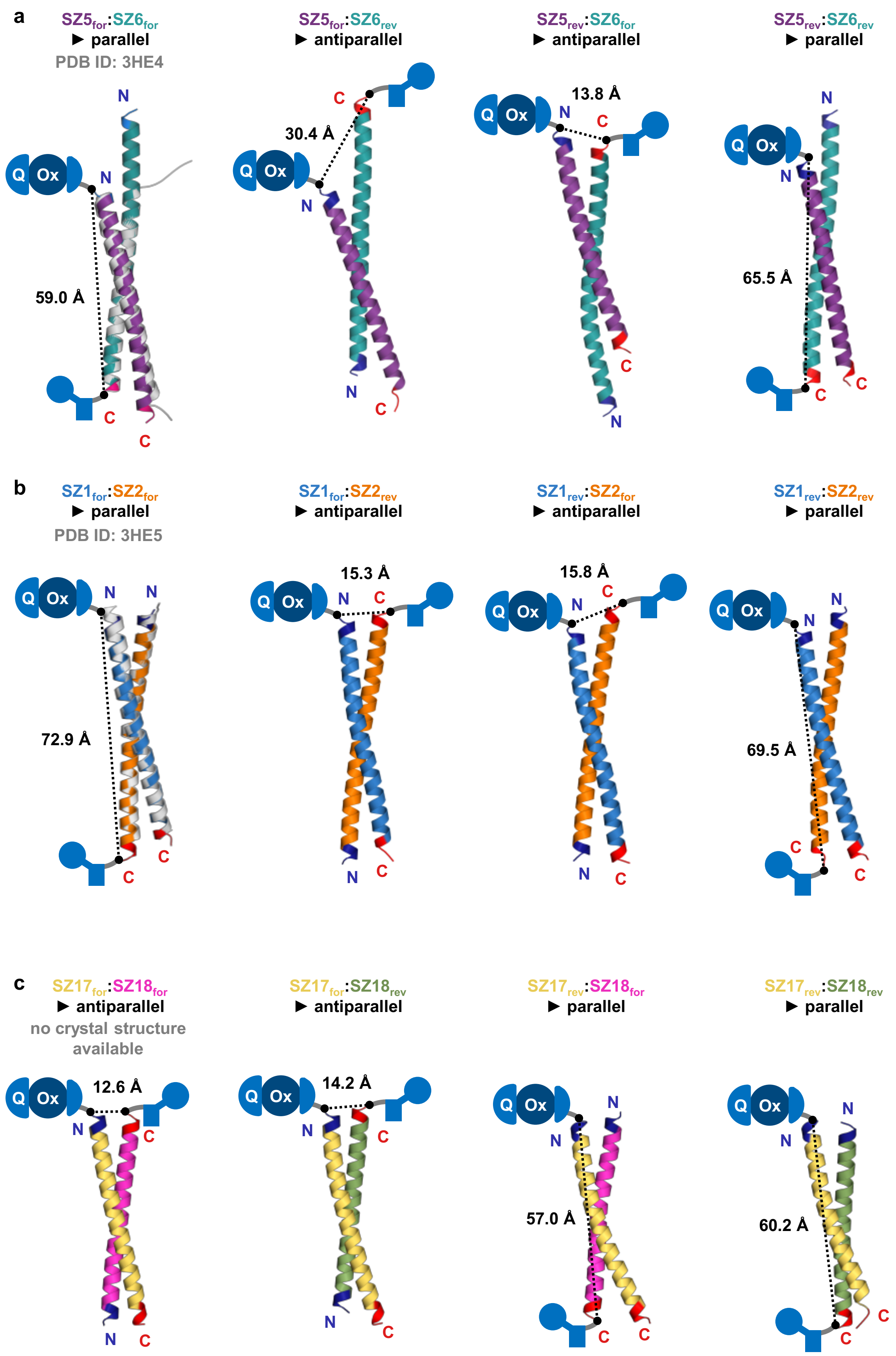
a
SZ5for:SZ6for
▶ parallel
PDB ID: 3HE4
59.0 Å
SZ5for:SZ6rev
▶ antiparallel
30.4 Å
SZ5rev:SZ6for
▶ antiparallel
13.8 Å
SZ5rev:SZ6rev
▶ parallel
65.5 Å
b
SZ1for:SZ2for
▶ parallel
PDB ID: 3HE5
72.9 Å
SZ1for:SZ2rev
▶ antiparallel
15.3 Å
SZ1rev:SZ2for
▶ antiparallel
15.8 Å
SZ1rev:SZ2rev
▶ parallel
69.5 Å
c
SZ17for:SZ18for
▶ antiparallel
no crystal structure available
12.6 Å
SZ17for:SZ18rev
▶ antiparallel
14.2 Å
SZ17rev:SZ18for
▶ parallel
57.0 Å
SZ17rev:SZ18rev
▶ parallel
60.2 Å
Q
Ox
N
C

**Figure S4| SYNZIP structure models and orientation**. All structural models were generated using AlphaFold 3. The predicted interaction geometry (parallel or antiparallel) and the predicted interdomain distance at the XUT[I] split site are depicted for each configuration. (**a**) All four orientation combinations of SZ5:SZ6: SZ5for:SZ6for (parallel), $SZ5_{for}$:$SZ6_{rev}$ (antiparallel), $SZ5_{rev}$:$SZ6_{for}$ (antiparallel), $SZ5_{rev}$:$SZ6_{rev}$ (parallel). The crystal structure of $SZ5_{for}$:$SZ6_{for}$ (PDB: 3HE4 displayed in grey) is shown for comparison. (**b**) All four orientation combinations of SZ1:SZ2: $SZ1_{for}$:$SZ2_{for}$ (parallel), $SZ1_{for}$:$SZ2_{rev}$; $SZ1_{rev}$:$SZ2_{for}$ (antiparallel), $SZ1_{rev}$:$SZ2_{rev}$ (parallel). The crystal structure of $SZ1_{for}$:$SZ2_{for}$ (PDB: 3HE5 displayed in grey) closely matches the AlphaFold3 prediction. (**c**) All four orientation combinations of SZ17:SZ18: $SZ17_{for}$:$SZ18_{for}$ (antiparallel), $SZ17_{for}$:$SZ18_{rev}$ (antiparallel), $SZ17_{rev}$:$SZ18_{for}$ (parallel), $SZ17_{rev}$:$SZ18_{rev}$ (parallel).
